# Drought disrupts volatile-mediated predator foraging and oviposition, weakening trait-mediated top-down control

**DOI:** 10.64898/2026.04.16.718996

**Authors:** Bijay Subedi, Jared Gregory Ali, Mônica F. Kersch-Becker

## Abstract

1. Drought is a major abiotic stressor that can restructure trophic interactions by limiting herbivore success and disrupting chemical signaling between plants and natural enemies. In tritrophic systems, plant volatiles guide natural enemy foraging and reproductive investment, often scaling with herbivore density; however, it is unclear whether drought alters this relationship and weakens top-down control.
2. Using a tomato-aphid-ladybeetle system, we tested how drought and herbivore density jointly affect plant VOC emissions, predator behavior, and aphid dynamics. We manipulated water availability (well-watered vs. drought) and aphid density (low vs. high), and measured plant physiology, volatile profiles, predator visitation and oviposition, and aphid responses.
3. Drought reduced stomatal conductance, plant biomass, and both total and compositional output of VOCs. Emission of key predator-attracting compounds (e.g., methyl salicylate, β-myrcene) peaked in well-watered, high-density plants but was suppressed under drought.
4. Ladybeetle visitation increased with aphid density but declined under drought, reflecting conserved shifts in volatile cues. Oviposition was concentrated on well-watered, high-density plants and associated with specific compounds (e.g., methyl salicylate, carvacrol), while others (e.g., cymene-7-ol, para, 1-octanol) were negatively associated.
5. Aphid suppression by predators occurred only under well-watered, high-density conditions. Under drought, aphid growth was already constrained, and predators had little additional effect on their abundance. However, both drought and predator presence influenced aphid demography, increasing production of dispersive alates.
6. These findings underscore the sensitivity of chemically mediated trophic interactions to environmental stress. Increased drought disrupts plant signaling, reducing natural enemy effectiveness, weakening biocontrol, and shifting herbivore population structure. Understanding how stress alters cue reliability is key to predicting community dynamics and managing ecosystem functions under stress.

## 1. Introduction

Natural enemies play an important role in regulating herbivore populations, often producing cascading effects on plant communities (Schmitz et al., 2000). However, the strength of top-down control often declines under abiotic stress (Barton & Schmitz, 2009; Lin et al., 2023). This weakening does not always result from natural enemy loss or herbivore escape (Clavijo McCormick, 2016; Schmitz & Barton, 2014). Observed declines in natural enemy efficacy have been linked to reductions in foraging, prey encounter rates, and reproductive investment; behavioral shifts that collectively undermine herbivore suppression despite apparent trophic continuity (Barton & Schmitz, 2009; Lin et al., 2022). These findings suggest that abiotic stress may disrupt not just species presence but also the behavioral mechanisms that sustain effective natural enemy-prey interactions. One emerging hypothesis is that stress degrades the informational channels, such as chemical cues, that natural enemies rely on to locate, evaluate, and commit to prey-rich habitats, severing top-down control through a breakdown in ecological communication rather than demographic loss (Holopainen et al., 2025; Pinto-Zevallos & Blande, 2024).

In plant-insect systems, natural enemies locate prey using plant volatile organic compounds (VOCs), including herbivore-induced plant volatiles (HIPVs), which are released in response to herbivory (Clavijo McCormick, 2016). HIPVs often convey information about prey presence, abundance, and quality, guiding natural enemy foraging, patch choice, and oviposition (Shiojiri et al., 2010; Turlings & Erb, 2018). However, volatile production can be metabolically costly and are tightly regulated by plant physiological status (Holopainen & Gershenzon, 2010). Drought stress disrupts hormonal signaling pathways, such as abscisic acid, jasmonic acid, and salicylic acid, that govern VOC biosynthesis (He et al., 2025; Holopainen & Gershenzon, 2010; Weldegergis et al., 2015). Such stress-driven disruptions may alter the composition, timing, or intensity of VOC emissions, thereby diminishing the reliability of these cues for predators, even when herbivore damage persists.

Many natural enemies respond to specific compounds or ratios within a blend, and slight changes in composition can disrupt recognition (Turlings & Erb, 2018). Changes in both major or minor components can compromise signal identity (Bruce et al., 2010), leading natural enemies to reduce searching, abandon prey-rich patches, or refrain from oviposition (Ali et al., 2023; De Rijk et al., 2016). Such shifts may reduce top-down control independently of natural enemy abundance. More broadly, natural enemy responses are likely shaped by the odor environment available when a patch is first encountered, which reflects an integrated plant response to ongoing herbivory rather than a momentary snapshot of damage. Oviposition is particularly sensitive: for many carnivorous insect, including ladybugs, VOC cues guide reproductive investment (Riddick, 2020; Verheggen et al., 2008; Xiu et al., 2019). Plant VOCs help them assess whether a site has sufficient prey to support offspring development (Peñaflor et al., 2011; Riddick, 2020). If drought reduces plant VOC reliability, natural enemies may forego oviposition, weakening top-down control across generations.

While drought often suppresses VOC production, some studies suggest that high herbivore densities may partially compensate by intensifying feeding damage, potentially enhancing HIPV output or shifting blend composition (Horiuchi et al., 2003; Shiojiri et al., 2010). This raises the possibility that increased herbivore density, by intensifying feeding damage, could, in theory, mitigate signal suppression under stress. However, the outcome likely depends on whether plants under stress maintain the capacity to scale plant VOC output with damage, and whether the resulting blends remain behaviorally relevant to natural enemies. In some cases, stress may favor deterrent or non-informative compounds that are less effective in guiding natural enemies (Rahman et al., 2025). Natural enemies often rely on a limited subset of volatiles, so-called “*keystone infochemicals*”, to make foraging and oviposition decisions (Ali et al., 2023; Turlings & Erb, 2018). This raises a critical question: does drought selectively reduce these key compounds, even when total emission or herbivore pressure remains high? If so, top-down control may fail not due to signal loss, but loss of important signaling compounds.

Understanding how stress shapes keystone volatiles is therefore essential for evaluating the resilience of chemical communication under environmental stressors.

Despite growing evidence that both abiotic and biotic factors shape plant VOC signaling, few studies test how both abiotic and biotic stressors interact to shape chemical signaling, predator behavior, and ecological outcomes. Most examine emission or behavior in isolation, limiting insight into full trophic pathway from cue production to herbivore suppression.

Crucially, the potential for herbivore density to modulate signal output under stress is rarely examined in tandem with predator decision-making. Even fewer address reproductive decisions, like oviposition, which may offer strong indicators of signal perception and predator commitment. Moreover, volatile blends are often treated as uniform signals, overlooking predators reliance on a limited subset of behaviorally active compounds. As a result, it is unknown whether density-driven amplification of plant VOCs under stress preserves the functional components needed for predator engagement, or if abiotic stress selectively undermines chemical communication even at high herbivore pressure. Understanding this interaction is key to predicting whether prey density can buffer, or is ultimately overridden by, abiotic constraints on top-down control.

Here, we test whether drought and herbivore density jointly influence trophic interactions by altering plant VOC signaling and predator responses. Specifically, we test whether: (1) drought reduces or alters plant VOC emission, (2) high herbivore density can restore signal quality or relevance, (3) predators adjust foraging and oviposition in response to these cues (4) specific plant VOCs are predictive of predator behavior, reflecting their role as ecologically relevant cues that guide predator foraging and oviposition, and whether these compounds are selectively suppressed under drought and (5) herbivore suppression by predator depends on both signal reliability and prey availability. By reframing predator-prey interactions as signal-contingent processes, this study proposes a new hypothesis in multitrophic ecology: one where the reliability, composition, and behavioral interpretability of cues determine the strength of top-down ecological regulation under stress.

## 2. Materials and methods

### 2.1 Study system

The tri-trophic model system comprised tomato (*Solanum lycopersicum* cv. Moneymaker) as the host plant, the potato aphid (*Macrosiphum euphorbiae* (Homoptera: Aphididae)) as the herbivore, and the convergent lady beetle (*Hippodamia convergens* (Coleoptera: Coccinellidae)) as a generalist predator. We conducted a greenhouse experiment between May and August 2025 to investigate how drought and herbivore density interact to influence plant physiology, volatile emissions, predator behavior, and trophic dynamics in a tritrophic system.

### 2.2 Plant cultivation and water treatments

Tomato seeds were sown in 10-cm pots containing Sunshine Mix #1 (Sungro Horticulture, USA) and maintained in a greenhouse under controlled environmental conditions (22 ± 2 °C, 16:8 h light:dark photoperiod) (Fig. 1). After germination, seedlings were thinned to one per pot and fertilized using a 15-9-12 NPK slow-release formulation (Osmocote Plus, Scotts Miracle-Gro).

**Fig. 1.**
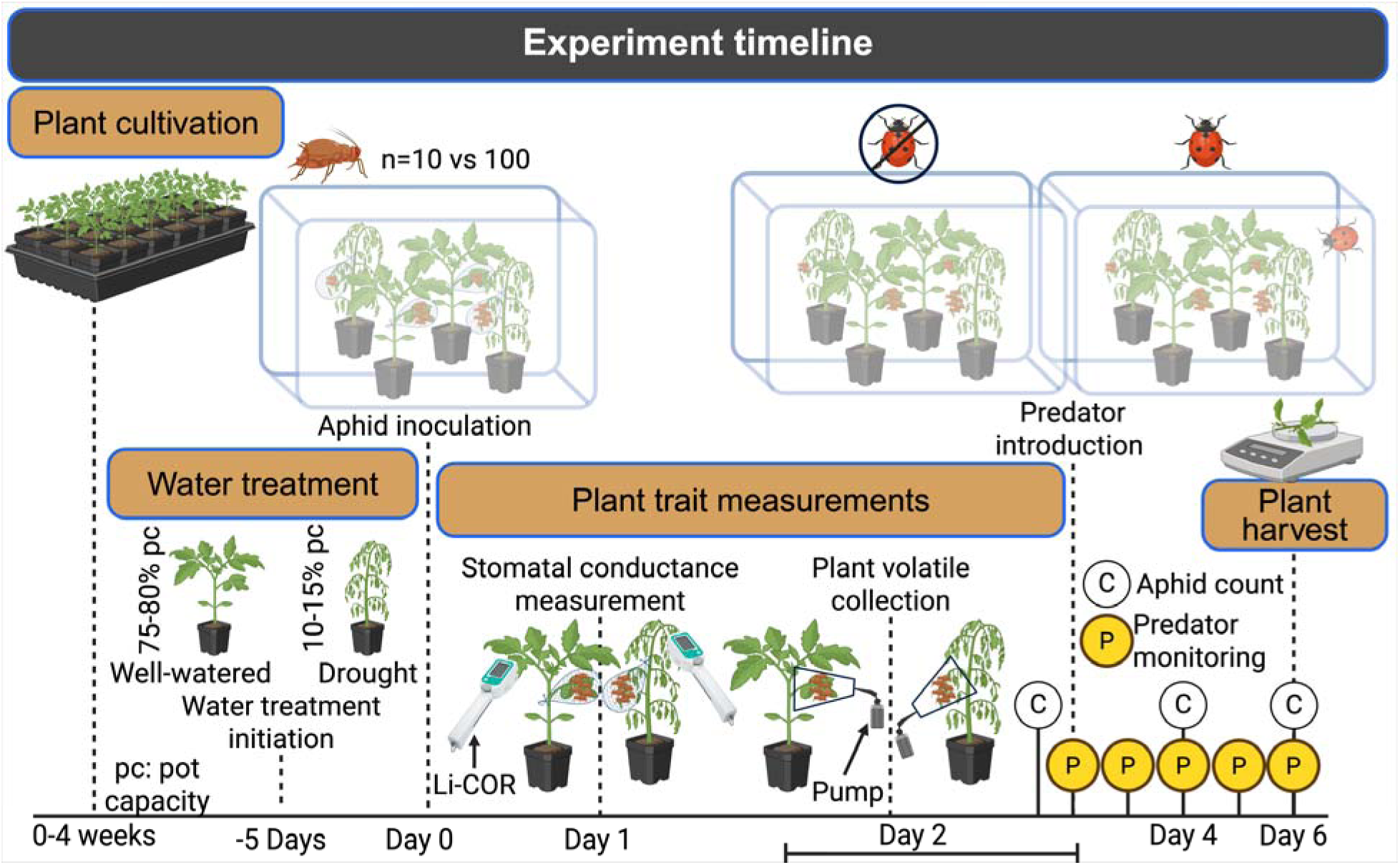
Experimental timeline and schematic overview of the experimental sequence showing key phases. The timeline illustrates when each major activity occurred, including plant trait measurements (stomatal conductance and volatile collection), aphid and predator introductions, behavioral observations, and final harvest for plant biomass.

Plants were allowed to grow for four weeks until they reached the four-leaf stage, at which point water treatments were initiated (Fig. 1).

To impose drought conditions, we manipulated soil volumetric moisture content beginning five days prior to aphid introduction. Well-watered plants were maintained at 75-80% of pot capacity, while drought-stressed plants were maintained at 10-15% (Fig. 1). We verified and adjusted moisture levels every other day using an ECOWITT WH0291 soil moisture probe (ECOWITT, Shenzhen, China), and calculated irrigation volumes using a gravimetric calibration curve.

### 2.3 Insect colonies and inoculation protocol

We maintained *M. euphorbiae* colonies on tomato plants in growth chambers set to 22 °C with a 16:8 h light:dark cycle. For experimental inoculations, we used a mixture of apterous adults and late-instar nymphs. Each plant received either 10 (Low) or 100 (High) apterous aphids of mixed age on its 3^rd^ fully expanded leaf, which was enclosed with fine mesh sleeve to allow the plants to emit HIPVs reflective of treatment intensity.

Predators were adult *H. convergens* beetles, which were reared on tomato plants with potato aphids at 23 °C with a 16:8 h light:dark cycle. Prior to release, we selected newly emerged, mated females for use in assays. To standardize foraging motivation, we starved individuals for 24 hours before introducing them into the experimental cages.

### 2.4 Greenhouse mesocosm setup

To assess predator foraging and oviposition, and aphid suppression across treatments, we established mesh cages mesocosms (40 × 60 × 100 cm; BioQuip Amber Lumite) in the greenhouse. The experiment followed a fully factorial design, manipulating two independent variables at plant level: water (well-watered vs. drought) and aphid density (Low (10 aphids per plant) vs. High (100 aphids per plant); Fig. 1). These treatments were fully crossed, resulting in four unique plant-level treatment combinations: well-watered with low aphid density, well-watered with high aphid density, drought-stressed with low aphid density, and drought-stressed with high aphid density. Each mesocosm (cage) contained one plant from each of these four treatment groups, arranged in randomized positions. A third factor, predator presence (present vs. absent), was manipulated at the cage level, with replicate cages assigned to either predator-present or predator-absent treatments (Fig. 1).

### 2.5 Plant traits

We measured stomatal conductance (*gsw*) using a LI-600 Porometer (LI-COR Environmental, Lincoln, NE, USA) on the fourth fully expanded leaf of each plant prior plant volatile collection (N = 14 replicates across two trials) (Fig. 1). At the end of the experiment, aboveground fresh biomass was quantified by harvesting and immediately weighing all shoot tissue using a precision balance (N = 14 replicates across two trials) (Fig. 1).

### 2.6 Volatile collection and chemical analysis

We collected plant VOCs 72 hours after aphid inoculation to test how aphid-density and water treatments shaped volatile emissions prior to predator introduction. Our VOC analysis was designed to compare treatment-specific odor environments, rather than to partition constitutive and aphid-induced emissions. We enclosed the third fully expanded leaf of each plant in a 1.2 L polyethylene cup and collected volatiles for six hours (airflow rate: 500 mL/min) using a custom-built headspace sampling system (Figure 4-1). Volatile compounds were trapped on HayeSep-Q filters (Supelco, Bellefonte, PA, USA), eluted with 200 μL of dichloromethane, and spiked with nonyl acetate (2 ng/μL) as an internal standard. Volatile extracts were analyzed using gas chromatography-mass spectrometry (GC-MS; Agilent 7890A/5975C). One microliter of each sample was injected in splitless mode at 250 °C, with helium as the carrier gas at 0.7 mL min ^1^. Compounds were separated on an HP-5MS column (30 m × 0.32 mm, 0.25 µm film thickness; Agilent Technologies) using an oven program of 40 °C for 2 min, followed by an increase of 10°C min ^1^ to 300 °C, with a final hold of 4 min. The MS was operated in positive EI mode. Peaks were deconvoluted in MassHunter Unknowns Analysis (Agilent Technologies, Santa Clara, CA, USA), tentatively identified by comparing mass spectra with the NIST17 and Adams libraries spectra and confirmed by matching retention index with published sources, including The Pherobase (Adams, 2007; Trase et al., 2025). The relative amounts of the detected compounds were then determined by referencing their total ion chromatogram (TIC) peak areas to those obtained for the internal standard, nonanyl acetate, at 2 ng/µL.

### 2.7 Predator behavior and aphid suppression

Following plant volatile collection, we released a single 24h-starved *H. convergens* female adult at the center of each cage (Fig. 1). Each treatment had 14 replicates across two trials. To evaluate sustained predator orientation over time, we monitored ladybeetle location in cage across six consecutive days. We recorded the plant occupied, or visitation behavior, of each individual at multiple timepoints across the trial period. Observations were made twice on days 2, 4, and 6, and three times on days 3 and 5, for a total of twelve observations per individual. In both trials, ladybug began ovipositing by Day 3. We recorded the presence and location of eggs at 72-, 88-, and 110-hours post-release to determine where ladybeetle chose to oviposit (i.e., oviposition behavior).

To assess predator-mediated aphid suppression, we conducted aphid counts at day 2, day 4, and day 6 after *H. convergens* release (Fig. 1). At each time point, we recorded the total number of aphids per plant, including both apterous and alate morphs, and counted the number of newly produced nymphs. Because aphids were free to move among plants within each cage, predator effects on individual plants could reflect a combination of consumptive (lethal) and non-consumptive (behavioral or trait-mediated) effects, including deterrence or spatial displacement.

### 2.9 Statistical analysis

All data were analyzed in R (v4.3.1).

*Plant traits:* Stomatal conductance was measured prior to predator introduction and thus was modeled as a function of water availability and aphid density (*water × density*). In contrast, fresh biomass was an endpoint measurement collected after predator exposure, and its model included all main effects and their interactions (*water × density × predator*). Each trait was analyzed using a generalized linear mixed model (GLMM) fitted with the *glmmTMB* package (Brooks et al., 2017). Fixed effects included all main effects and their interactions, while plant ID nested within cage and block (1| Block/Cage/Plant ID) was included as a random effect to account for non-independence within experimental units. For stomatal conductance, we specified a *Tweedie* distribution to account for overdispersion and non-Gaussian error structure. For fresh biomass, a *Gaussian* distribution was appropriate based on residual diagnostics. Model fit and dispersion were assessed using simulated residuals via the *DHARMa* package (Hartig & Hartig, 2017).

Type III Wald χ^2^ tests were conducted using the *Anova*() function from the *car* package (Fox & Weisberg, 2018) to evaluate the significance of fixed effects. We performed post-hoc comparisons using *estimated marginal means* (*emmeans* package) (Lenth & Lenth, 2018), and visualized group differences with compact letter displays using *Tukey*-adjusted comparisons via the *multcomp* (Hothorn et al., 2016).

*Plant volatile organic compounds (VOCs):* To examine plant volatile profiles in response to aphid density and water availability, we collected VOCs prior to predator introduction (see Section 2.6). We analyzed VOC data across two complementary approaches:

*(a) Volatile composition:* We evaluated multivariate differences in plant VOC composition using permutational multivariate analysis of variance (PERMANOVA) via the adonis2() function in the vegan package (Oksanen et al., 2013). The analysis was based on Jaccard distances calculated from presence/absence-transformed VOC data with 999 permutations, testing the effects of water treatment, aphid density, and their interaction. We tested the effects of water treatment, aphid density, and their interaction, and we assessed homogeneity of dispersion with *betadisper*(). We further explored significant treatment contrasts with pairwise PERMANOVA and applied FDR correction to the resulting P-values. To visualize VOC composition patterns across treatments, we used CAP/dbRDA (capscale()), and to identify individual VOCs associated with treatment separation, we fitted compounds to the CAP ordination with envfit(), using 999 permutations and FDR correction across compounds.
*(b) Volatile analysis by individual compounds:* A multivariate analysis of variance (MANOVA) was not used due to violations of its core assumptions. All response variables (VOCs) deviated significantly from normality (Shapiro-Wilk P<0.001). Given the zero-inflated and non-normal nature of VOC data, we analyzed the emission rates of individual plant VOCs using GLMMs (glmmTMB package) with tweedie distribution and zero inflation. Fixed effects included water treatment (well-watered vs. drought), aphid density (Low (10) vs. high (100)), and their interaction, with plant ID nested within cage and block included as a random effect. Type II Wald χ^2^ tests (car package) were used to assess fixed effects.

*Predator behavioral responses*: To capture the full spectrum of ladybeetle behavioral responses, we conducted separate analyses for two distinct behaviors: visitation (i.e., plant choice) and oviposition. These behaviors were modeled independently using statistical frameworks appropriate for each response type, as described below.

*Predator visitation behavior:* We analyzed the ladybeetle visitation behavior (i.e., visits or choices) using a conditional logit model (*clogit*, *survival* package (Therneau, 2015)) with plant visitation (chosen = 1 vs. 0) as the binary response. We included water treatment (well-watered vs. drought), inoculation density (low (10) vs. high (100)), and their interaction as fixed effects, and stratified model by choice set (*cage* × *timepoint*) to account for repeated measures within cages. We assessed the significance of model terms using Type III Wald χ^2^ tests (*car* package).

*Predator oviposition behavior*: We analyzed ladybeetle oviposition behavior using a Bayesian logistic regression framework implemented in the *brms* package (Bürkner, 2017). The binary response variable indicated whether or not an individual plant received any oviposition (1 = oviposition, 0 = none). Each plant was classified by its water availability treatment (well-watered or drought) and aphid density (low (10) or high (100)). We modeled these effects across three time points (72, 88, and 110 hours after introduction) and incorporated a random intercept for each unique choice set (defined by trial, cage, and time point) to account for non-independence within trials. The primary model included fixed effects for time point, water treatment, aphid density, and their two-way interactions (*time × water, time × density*). This model was preferred over the full three-way interaction model (*time × water × density*), which showed poor convergence (high R-hat and exceeded tree depth), likely due to overparameterization relative to sample size. A second model, collapsed across time points, included only *water × density* and a random intercept for choice set (i.e., cage). All models used a Bernoulli distribution with a logit link function. We ran four chains of 4000 iterations each (2000 warmup), using *adapt*_*delta* = 0.99 and *max_treedepth* = 12 for robust sampling. Posterior summaries were derived using median estimates and 95% highest posterior density (HPD) credible intervals.

*Role of plant volatiles in ladybeetle visitation and oviposition:* We evaluated how plant volatiles influenced two binary behavioral responses by the ladybeetle *H. convergens*: (1) visitation and (2) oviposition. We modeled predator visitation and oviposition as a function of VOC profiles measured following sustained herbivory. Because herbivore-induced VOC blends reflect integrated plant responses to herbivory and provide a biologically relevant snapshot of the odor environment available to predators at the onset of the assay, these profiles were used as proxies for the cues underlying predator decision-making despite non-synchronous sampling. In both cases, the response variable was coded as 1 if the event occurred (i.e., ≥1 choice or egg laid) or 0 otherwise. To reduce dimensionality and identify informative predictors, we first used a random forest classifier (*randomForest* package) (Liaw & Wiener, 2002) to rank volatile compounds by importance. We retained the top 19 compounds that contributed most to classification accuracy (for oviposition) or node purity (for choice).

We then fit Bayesian logistic regression models using the *brms* package, specifying a Bernoulli likelihood with a logit link and weakly informative priors (normal(0, 5)) on the regression coefficients. Predictors were z-score standardized prior to model fitting. Model performance was assessed using Bayesian R^2^, posterior predictive checks, and leave-one-out cross-validation (*loo* package) (Vehtari et al., 2021). To further identify the most predictive volatiles, we applied projection predictive variable selection using the *projpred* package (Piironen et al., 2023). Final reduced models were refit using only the selected subset of volatiles. For each reduced model, we extracted posterior summaries using the *bayestestR* package (Makowski et al., 2019) and interpreted effects as significant when 95% credible intervals excluded zero. To evaluate binary classification performance, we used the *posterior epred()* function to compute posterior mean probabilities for each observation. Predictions had a threshold at 0.5, and classification metrics including accuracy, sensitivity, specificity, precision, and F1 score were calculated against the observed binary responses. Receiver operating characteristic (ROC) curves and area under the curve (AUC) values were generated using the *pROC* package (Robin et al., 2021).

*Predator suppression of aphid population and demographics:* To evaluate the effects of predator presence, water availability, and aphid density on aphid population dynamics, we analyzed four response variables: aphid population growth rate (calculated as: *dN/Ndt*) = ln (*N_2_*- *N_1_*)/(*t_2_*-*t_1_*), where *N_1_* and *N_2_* are initial and final aphid densities at time *t_2_* and *t_1_*, respectively (Gotelli, 1995)), final density, nymph count, and alate count. Each response was modeled separately using GLMMs implemented in the *glmmTMB* package. Growth rate was treated as a continuous variable and modeled using a Gaussian distribution. The count-based responses were modeled using a negative binomial distribution to account for overdispersion. Models initially included all main effects and interactions among water, inoculation density, predator, and day (representing the three time points: day 2, 4, and 6), with final models selected by comparing full and reduced versions based on AIC and model diagnostics. The variable ‘day’ was retained in all models to account for temporal structure, and its interactions with other predictors were considered when relevant. To account for the nested experimental design, we included plant ID nested within cage and block as a random effect in all models. Residual diagnostics were performed using the *DHARMa* package to assess model fit, dispersion, and distributional assumptions. Significance of main effects and interactions was assessed using Type III Wald chi-square tests via the *Anova*() function (*car* package). For post hoc comparisons, we estimated marginal means using the *emmeans* package and visualized group differences using compact letter displays (*multcomp* package) with Tukey-adjusted comparisons.

## 3. Results

### 3.1 Plant traits

Stomatal conductance was reduced by approximately 96% under drought conditions (χ^2^ =118.93, *P*<0.001), while aphid density (χ^2^ =0.09, *P*=0.76) and its interaction with water availability (χ^2^ =0.004, *P*=0.95; Fig. 2a) were not significant. Because stomatal conductance was measured prior to predator introduction, predator presence was not included in this model. For aboveground fresh biomass, drought also led to approximately 74% reduction (χ^2^ =249.32, *P*<0.001), while aphid density (χ^2^ =0.01, *P*=0.91), predator presence (χ^2^ =0.15, *P*=0.70), and their two-way interactions (water × density: χ^2^ =1.27, *P*=0.26; water × predator: χ^2^ =0.44, *P*=0.51; density × predator: χ^2^ =0.10, *P*=0.75; Fig. 2b) were non-significant. The three-way interaction among water availability, aphid density, and predator presence was also not significant (χ^2^ =0.004, *P*=0.95; Fig. 2b).

**Fig. 2.**
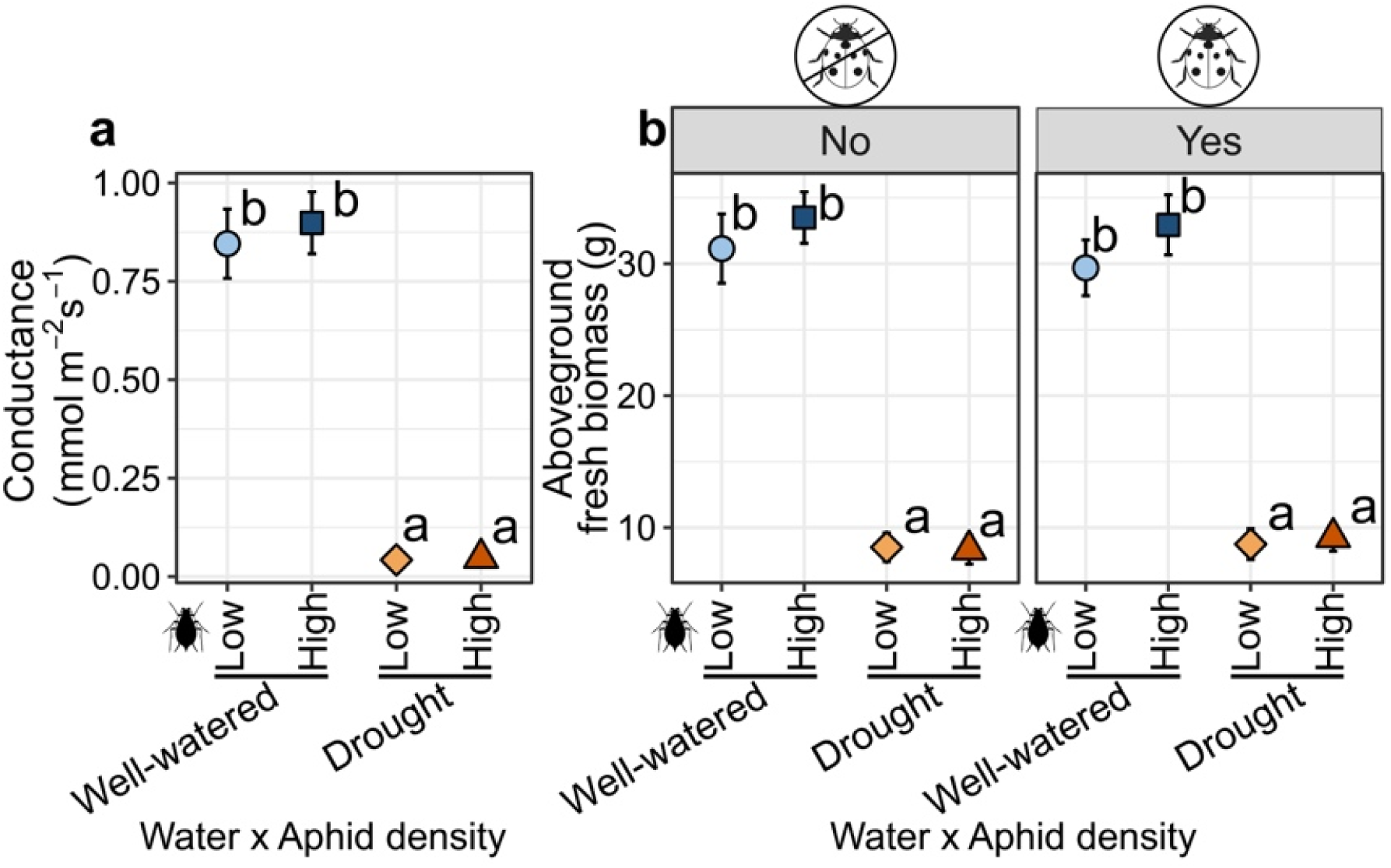
Effects of aphid density (low vs. high) and water availability (well-watered vs. drought) on (a) stomatal conductance (mmol m^-2^ s^-1^) and (b) aboveground fresh biomass (g). In (b), columns show predator presence (left: absent; right: present). Stomatal conductance was measured before predator introduction. Points show mean ± SE. Blue = well-watered; orange = drought; lighter shades = low aphid density, darker = high. Different letters indicate significant treatment differences (Tukey-adjusted, *P*<0.05).

### 3.2 Plant VOCs

*(a) Volatile composition:* PERMANOVA based on Jaccard dissimilarities of presence–absence volatile profiles revealed a significant water treatment × aphid density interaction (F_[1,106]_=0.96, R^2^=0.0087, *P*=0.042; Fig. 3), indicating that treatment combinations differed in VOC composition. Tests for homogeneity of multivariate dispersion were not significant (betadisper: F_[3,106]_=0.123, *P*=0.947; permutation test *P*=0.949), showing that differences among treatments were not driven by unequal within-group dispersion. Pairwise PERMANOVA comparisons revealed several significant differences in VOC profiles across specific treatment combinations. Pairwise PERMANOVA indicated significant differences in community composition among all treatment combinations after FDR correction. The strongest differences involving the well-watered low density treatment were observed relative to drought high density (F_[1,53]_=1.80, R^2^=0.0329, *P*=0.0012; Fig. 3), drought low density ((F_[1,53]_=1.67, R^2^=0.0306, *P*=0.0012), and well-watered high density (F_[1,52]_=1.17, R^2^=0.0220, *P*=0.0012). Well-watered high density also differed significantly from drought low density (F_[1,53]_=1.82, R^2^=0.0332, *P*=0.0012) and drought high density (F_[1,53]_=1.61, R^2^=0.0295, *P*=0.0012; Fig. 3). The contrast between drought low and drought high density was also significant, though weaker in magnitude than the other pairwise comparisons (F_[1,54]_=1.08, R^2^=0.0196, *P*=0.009). Environmental fitting *(envfit)* analysis of NMDS ordination identified 37 volatile compounds that were significantly associated with the ordination axes (*P*<0.05), indicating that their emission patterns were structured by treatment effects. Full results, including vector coordinates, R^2^ values, and significance levels, are presented in Supplementary table S1.
*(b) Volatile analysis by individual compounds:* Volatile composition differed among treatment combinations at the multivariate level. A principal components MANOVA on the first 20 PCs detected a significant Water × Density interaction (Pillai’s trace = 0.347, F_[20,87]_=2.31, *P*=0.004), demonstrating that the effect of water regime on the volatile blend depended on aphid density. To attribute this interaction to individual compounds, we combined PCA-derived interaction driver scores with compound-specific generalized linear mixed models. Driver scores quantify each compound’s contribution to the multivariate Water × Density separation, while mixed models provide inferential tests of Water, Density, and their interaction. All Wald χ^2^ statistics (df=1) and P values are presented in Table S2.

Eight compounds exhibited significant Water × Density interactions, directly supporting the non-additive multivariate response. An additional 20 compounds showed significant main effects in the absence of interaction: 13 exhibited a significant effect of Water only, 2 exhibited a significant effect of Density only, and 5 exhibited significant effects of both factors. In total, 28 compounds displayed at least one significant fixed effect and are reported in Table 1 and Supplementary Table S2.

**Fig. 3.**
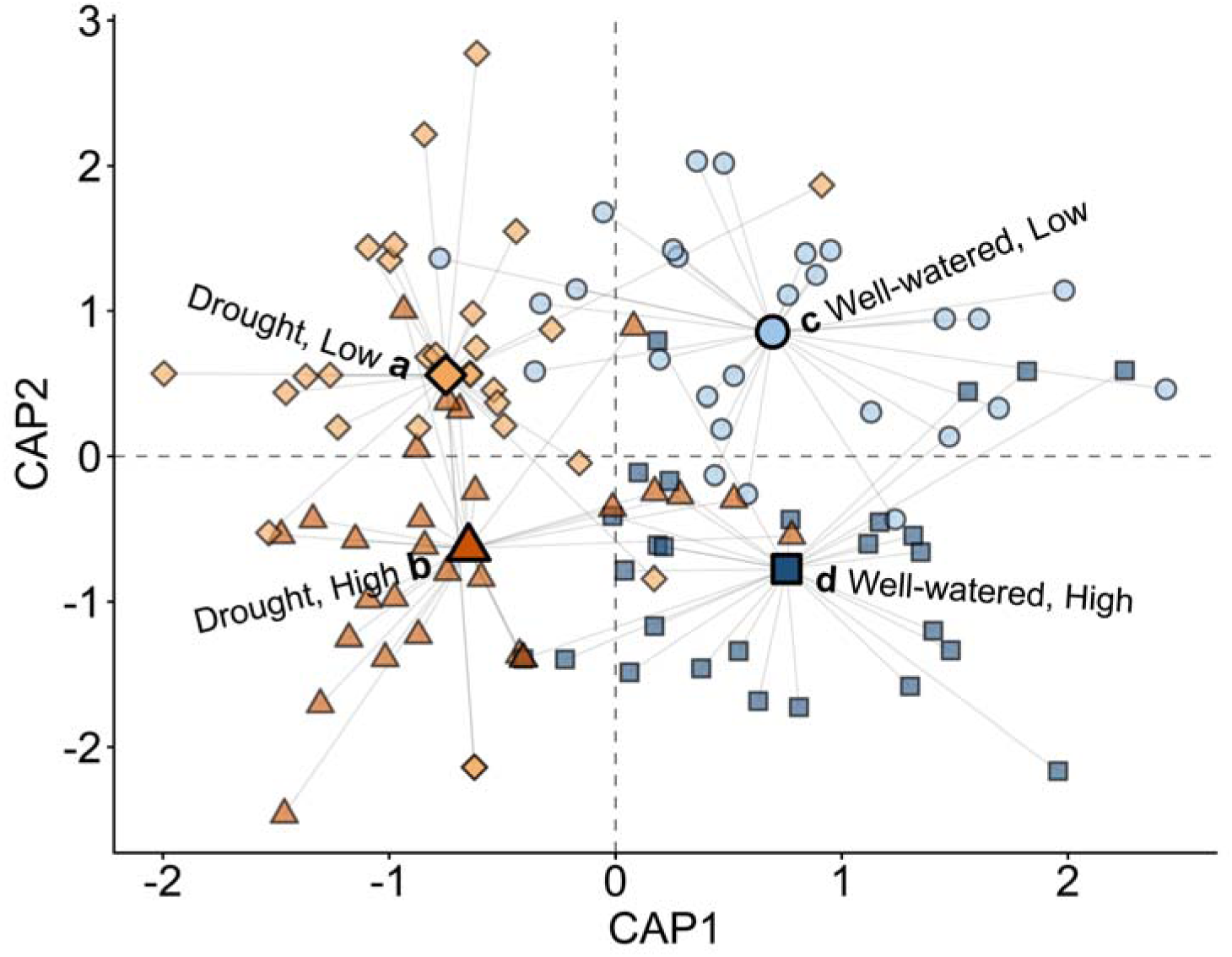
Constrained analysis of principal coordinates (CAP/dbRDA) of plant VOC composition based on Jaccard dissimilarities calculated from presence/absence data across water and aphid density treatments. Points represent individual plants, with orange indicating drought and blue indicating well-watered treatments. Shapes denote treatment combinations: low aphid density as circles or diamonds and high aphid density as squares or triangles. Large symbols indicate treatment centroids, and gray line segments connect individual samples to their respective centroids. Letters identify treatment groups for visual reference.

**Fig. 4.**
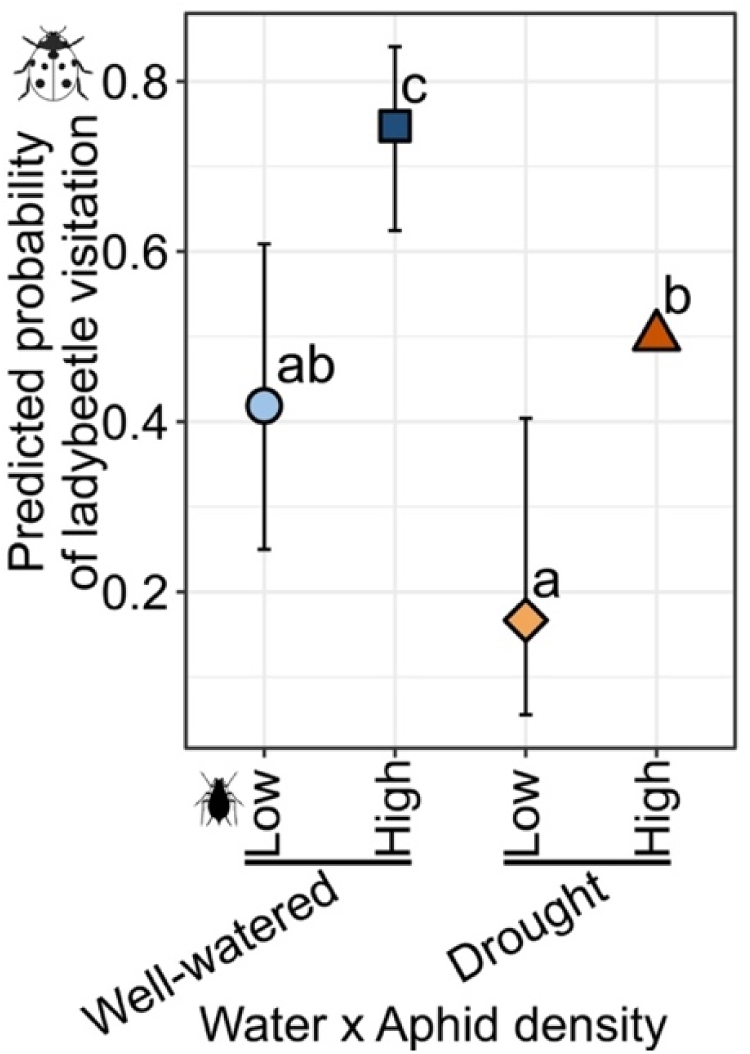
Predicted probability of predator visitation across aphid density and water availability treatments. Points show estimated marginal means (±95% CI) from a conditional logistic regression. Blue = well-watered; orange = drought; lighter = low aphid density, darker = high. Letters denote significant differences (Tukey-adjusted, P<0.05).

**Table 1:**
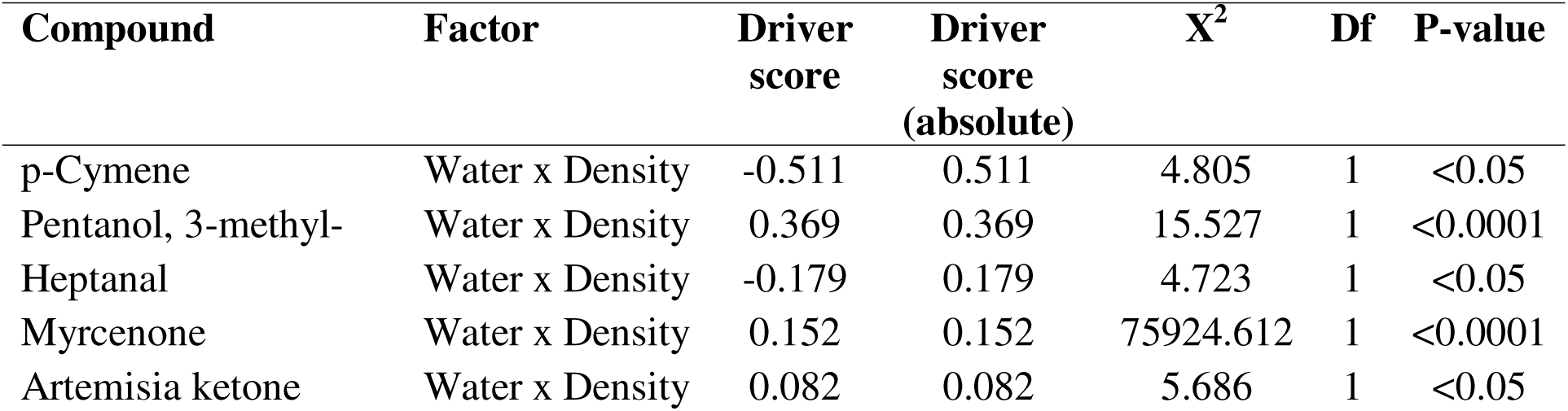
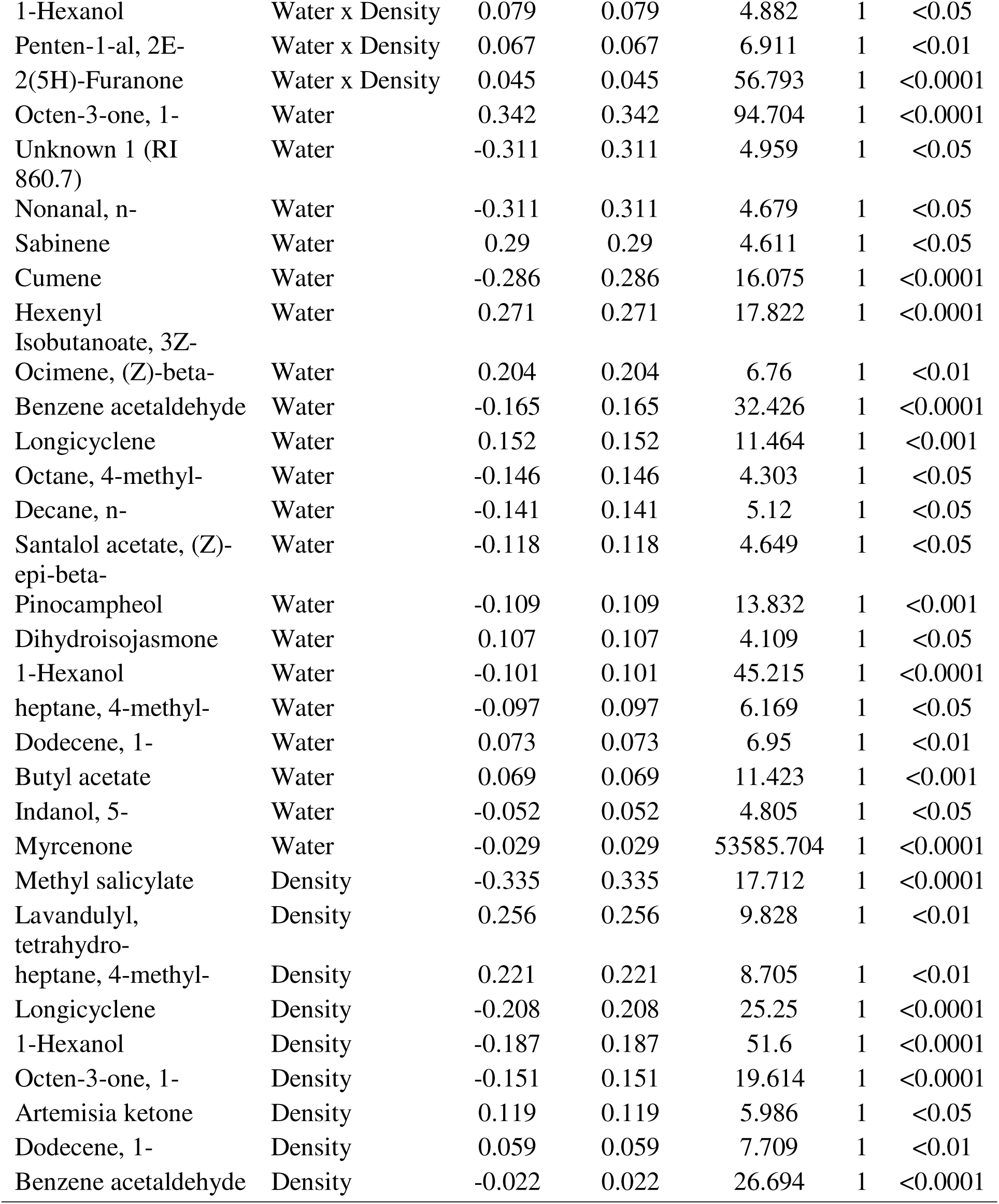
Compounds exhibiting significant fixed effects in generalized linear mixed models. For each compound and significant factor (Water × Density interaction, Water, or Density), the table reports the factor-specific multivariate driver score, its absolute magnitude (|driver score|), and the corresponding Wald χ^2^ statistic (Df = 1) with *P-*value. Driver scores were derived by projecting the principal components effect vectors for each factor back into compound space using PCA loadings and quantify each compound’s contribution to multivariate treatment separation. Only statistically significant effects (*P* < 0.05) are shown.

### 3.3 Predator preference and oviposition behavior

*(a) Predator visitation behavior:* Predator were approximately 4.5 times more likely to select plants with high-aphid density compared to those with low-aphid density (χ^2^ =10.79, *P*=0.001; Fig. 4). Drought stress reduced visitation, with ladybeetles 69% less likely to choose drought-stressed plants over well-watered ones (χ^2^ =22.01, *P*<0.001; Fig. 4). There was no significant interaction between aphid density and water availability (χ^2^ =0.12, *P*=0.725; Fig. 4), indicating that their effects were additive.
*(b) Predator oviposition behavior:* The probability of oviposition by female ladybeetle varied with plant water availability, aphid prey density, and time (Fig. 5). Across all time points, oviposition was highest on well-watered plants with high aphid density (probability = 0.477; 95% HPD: 0.331–0.635). In contrast, drought-stressed plants with low aphid density had near-zero oviposition (0.000; 95% HPD: 0.000–0.013). Intermediate probabilities occurred on well-watered low-aphid (0.160; 95% HPD: 0.064–0.282) and drought high-aphid plants (0.161; 95% HPD: 0.057–0.277), indicating independent and interactive effects of water and prey cues.

Temporal patterns revealed peak selectivity early in the experiment, before any eggs were present. At 72 hours, oviposition was strongly concentrated on well-watered/high-aphid plants (0.727; 95% HPD: 0.492–0.935), with little to none on other treatments. At 88 hours, this preference persisted (0.604; 95% HPD: 0.353–0.835), though oviposition on drought/high-aphid plants increased slightly (0.105; 95% HPD: 0.005–0.272). By 110 hours, oviposition probabilities converged across treatments (range: 0.092–0.242), with overlapping intervals suggesting reduced selectivity as eggs accumulated.

**Fig. 5.**
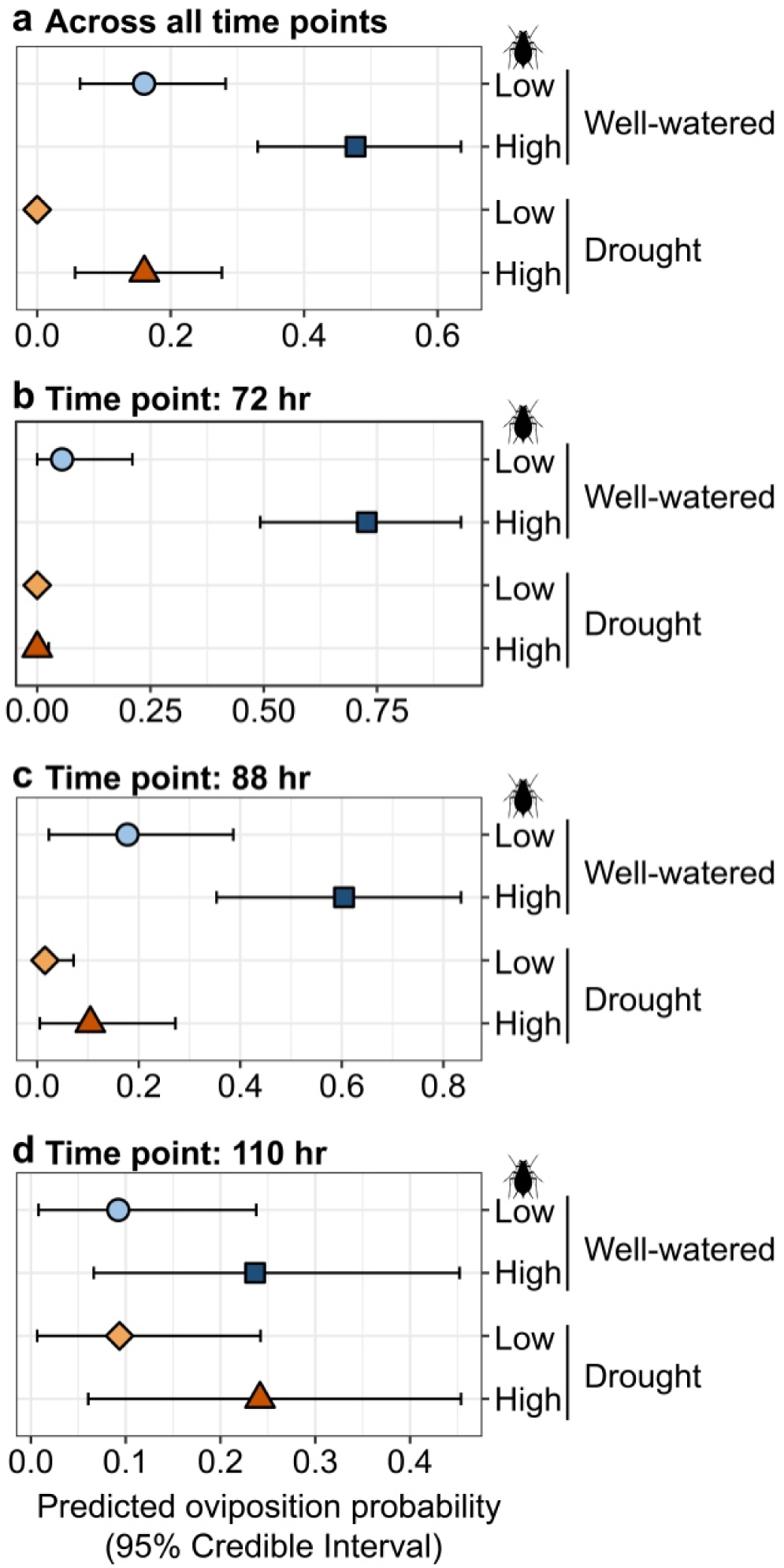
Predicted oviposition probabilities of female ladybeetles across water (well-watered vs. drought-stressed) and aphid density (low vs. high) treatments, shown cumulatively and by time point. Posterior means and 95% HPD intervals are based on a Bayesian logistic regression. Panel (a) shows estimates across all time points; panels (b)–(d) show predictions at 72, 88, and 110 hours. Circles/squares = well-watered (low/high density); diamonds/triangles = drought-stressed (low/high). Lighter shades = low, and darker = high aphid density.

Model estimates confirmed strong effects of aphid density and water status: oviposition was less likely on low-aphid plants (estimate = −19.40; 95% CI: −105.01 to −2.09), and more likely under well-watered conditions (estimate = 1.59; 95% CI: 0.55–2.71). A positive interaction between well-watered plants and low aphid density (estimate = 17.81; 95% CI: 0.36–103.67) suggests flexible decision-making based on cue combinations.

### 3.4 Predator visitation behavior in response to plant volatiles

Bayesian logistic regression identified four volatile compounds with credible associations with *H. convergens* olfactory choice behavior. The model included 19 candidate compounds selected from the random forest importance ranking and evaluated in a Bayesian framework. Overall model fit was moderate, with a Bayesian R-squared of 0.44 and a 95% credible interval of 0.33 to 0.52, and leave-one-out cross-validation indicated reasonable predictive performance with a LOOIC of 116.3. Four compounds had 95% credible intervals that excluded zero, indicating credible effects on ladybeetle choice probability (Fig. 6a-d). Three compounds were positively associated with the probability of ladybeetle choice: methyl salicylate (β = 4.51, 95% credible interval = 1.04 to 9.06; Fig. 6a), β-myrcene (β = 4.50, 95% credible interval = 1.30 to 8.09; Fig. 6b), and valencene (β = 2.77, 95% credible interval = 0.67 to 5.06; Fig. 6c). In contrast, 1-octanol was negatively associated with choice probability (β =-2.69, 95% credible interval =-5.41 to-0.17; Fig. 6d). Model discrimination was strong, with an area under the ROC curve of 0.903. Using a 0.5 classification threshold, the model achieved an accuracy of 0.833, sensitivity of 0.943, specificity of 0.538, precision of 0.846, and an F1 score of 0.892. Together, these results indicate that adult ladybeetles respond selectively to particular plant volatiles, with methyl salicylate, beta-myrcene, and valencene acting as potential attractant-associated cues, whereas 1-octanol appears to reduce the likelihood of choice.

**Fig. 6.**
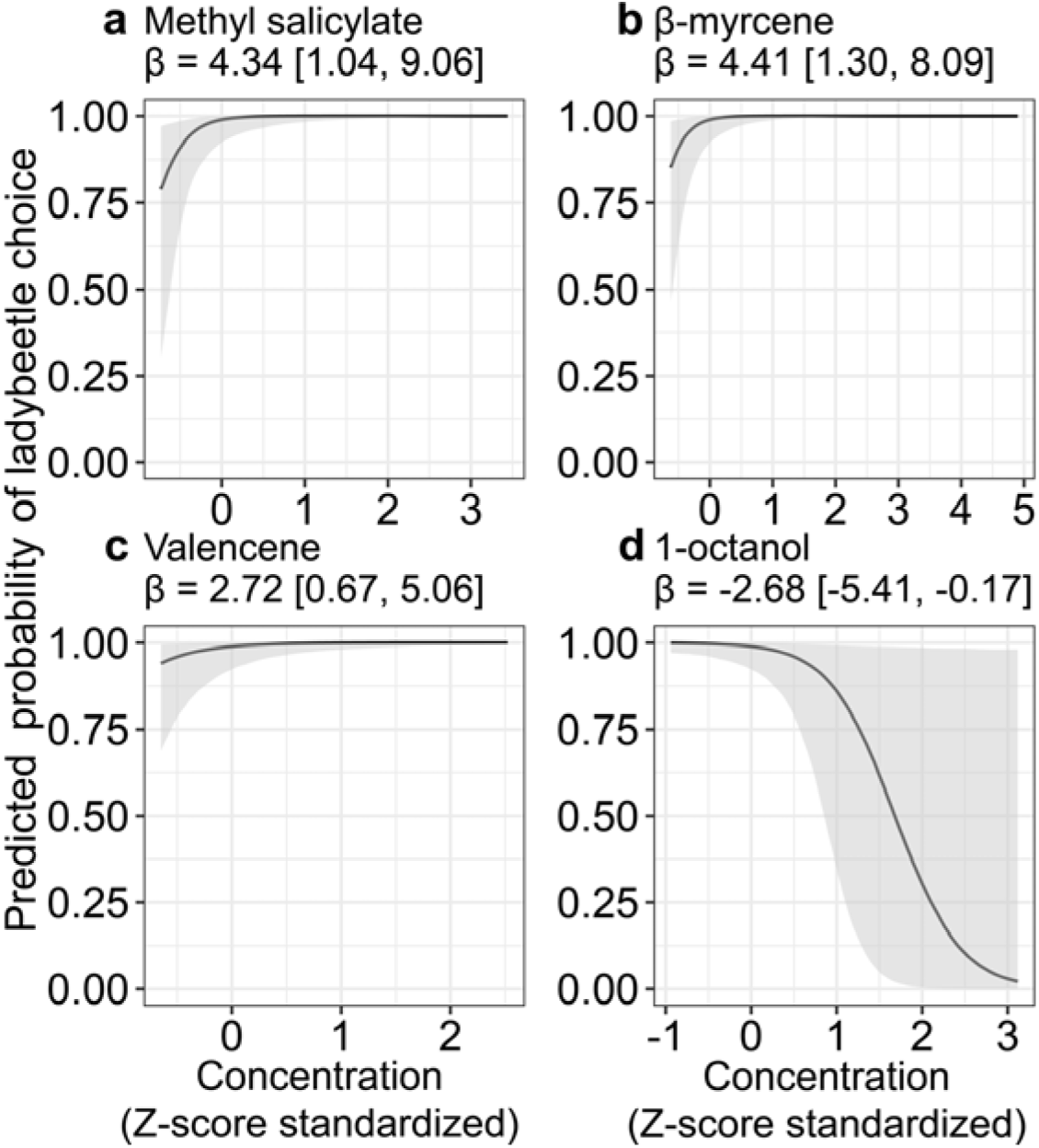
Predicted probability of ladybug preference from Bayesian logistic regression as a function of compound concentration (Z-score standardized). Shaded areas represent 95% credible intervals. Labels above each panel show the posterior median and 95% credible interval for the estimated effect (β).

### 3.4 Predator oviposition behavior in responses to volatile compounds

Bayesian logistic regression identified 12 volatile compounds with credible associations with *H. convergens* oviposition behavior. The reduced model included 19 candidate compounds and showed strong explanatory power, with a Bayesian R-squared of 0.69 and a 95% credible interval of 0.58 to 0.79. Leave-one-out cross-validation indicated good apparent predictive performance, with a LOOIC of 87.0, although several high Pareto k values suggest that these cross-validation results should be interpreted cautiously. Twelve compounds had 95% credible intervals that excluded zero, indicating credible effects on oviposition probability (Fig. 7a to i). Five compounds were positively associated with ladybeetle oviposition: methyl salicylate (β = 6.22, 95% credible interval = 3.08 to 10.08; Fig. 7a), n-heptanal (β = 6.13, 95% credible interval = 2.75 to 9.79; Fig. 7b), p-cymene (β = 6.31, 95% credible interval = 2.01 to 11.15; Fig. 7c), carvacrol (β = 4.64, 95% credible interval = 1.32 to 8.76; Fig. 7d), and 1-hexanol (β = 3.15, 95% credible interval = 0.98 to 5.49; Fig. 7e). Seven compounds were negatively associated with oviposition: 1-octanol (β =-3.29, 95% credible interval =-6.98 to-0.45; Fig. 7f), n-hexanal (beta =-6.15, 95% credible interval =-10.13 to-2.64; Fig. 7g), dihydroisojasmone (β =-5.39, 95% credible interval =-9.35 to-1.96; Fig. 7h), n-nonanal (β =-3.47, 95% credible interval =-6.58 to-0.62; Fig. 7i), α-pinene (β =-4.09, 95% credible interval =-7.40 to-0.98; Fig. 7j), cryptone (β =-5.83, 95% credible interval =-9.85 to-2.16; Fig. 7k), and para-cymen-7-ol (β =-3.36, 95% credible interval =-6.41 to-0.73; Fig. 7i). Model discrimination was extremely strong, with an area under the ROC curve of 0.990. Using a 0.5 classification threshold, the model achieved an accuracy of 0.958, sensitivity of 0.955, specificity of 0.962, precision of 0.955, and an F1 score of 0.955. Together, these findings indicate that adult ladybeetle oviposition potentially responds strongly and selectively to particular volatile compounds, with some compounds increasing the probability of egg laying and others acting as deterrent cues.

**Fig. 7.**
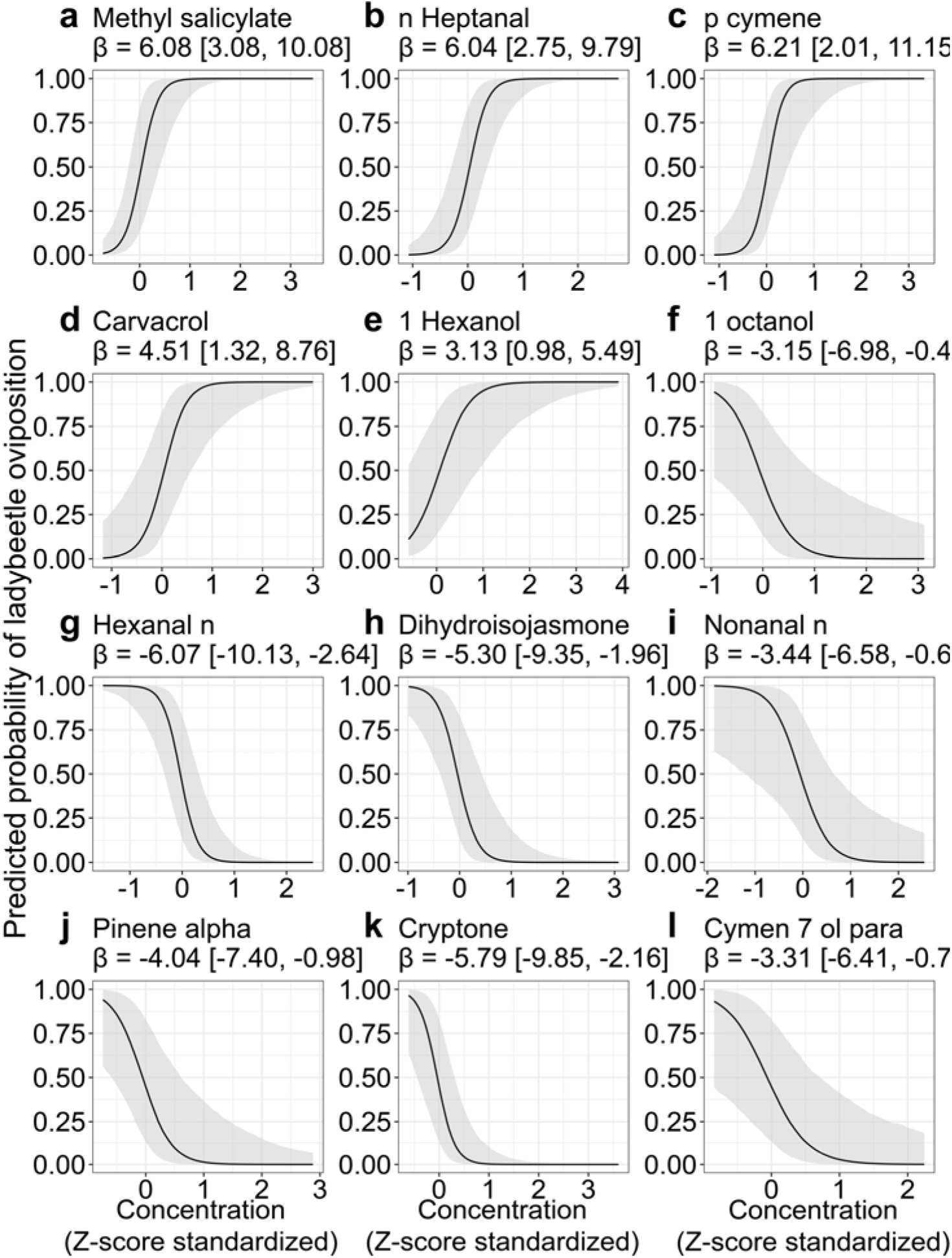
Predicted probability of oviposition from Bayesian logistic regression as a function of compound concentration (Z-score standardized). Each panel shows the marginal effect of a selected compound or treatment, with 95% credible intervals (shaded). Posterior coefficient (β) and its 95% CI are shown in panel titles.

### 3.6 Predator suppression of aphid population and demographics

Per capita aphid population growth rate varied significantly over time and was shaped by interactions among water availability and predator presence (*water × predator × day*: χ^2^ =6.85, *P*=0.032; Fig. 8a). Aphid inoculation density alone did not significantly affect per capita growth rates (χ^2^ =0.0629, *P*=0.802). Under well-watered and high aphid density, predator presence reduced aphid per capita population growth rate by approximately 70%, whereas under drought, growth remained near zero regardless of predator presence.

**Fig. 8.**
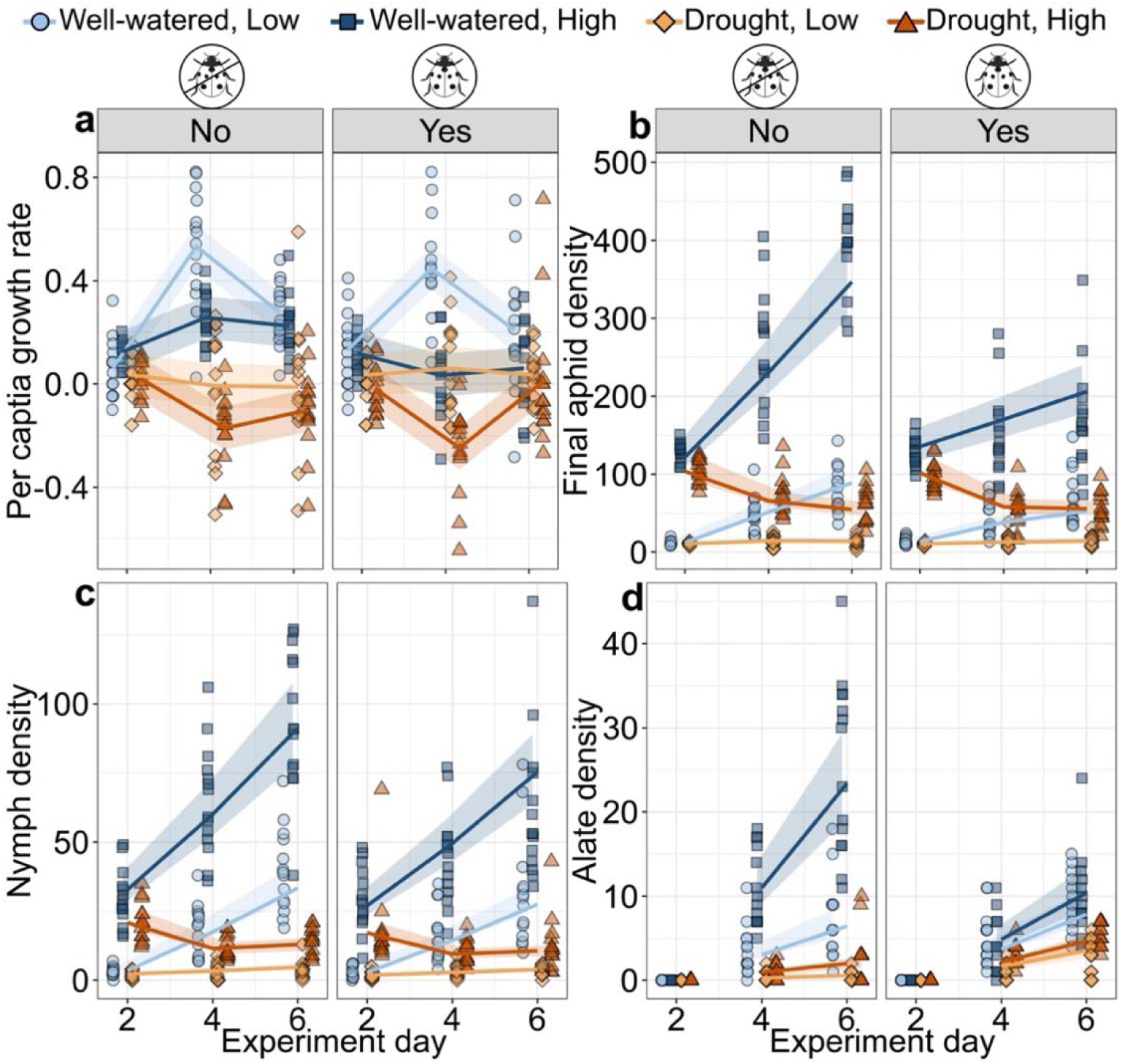
Effects of water availability, initial aphid density, and predator presence on aphid population growth and demography over time. Panels show (a) per capita growth rate, (b) final density, (c) nymph density, and (d) alate density under combinations of water (well-watered vs. drought), aphid density (low vs. high), and predator presence (No vs. Yes). Lines show model-predicted means ± 95% CI; points are individual replicates. Blue = well-watered; orange = drought; lighter = low aphid density, darker = high aphid density.

Final aphid density was driven largely by initial aphid density, time, and their interactions with water availability (*density × day:* χ^2^ =153.47, *P*<0.001; *water × day*: χ^2^ =248.43, *P*<0.001; Fig. 8b). A significant three-way interaction (*water × predator × day*: χ^2^ =19.11, *P*<0.001) confirmed that predator effects depended on both resource supply and time. Predator presence alone did not significantly affect overall aphid densities (χ^2^ =0.03, *P*=0.86). However, in well-watered, high-density treatments, predators reduced final aphid numbers by roughly 40% by the end of the experiment. Under drought, predator effects were negligible.

For nymph densities, significant interactions occurred for *water × day* (χ^2^ =163.50, *P*<0.001) and *day* × *density* (χ^2^ =91.28, *P*<0.001; Fig. 8c), indicating that nymph production was both temporally dynamic and resource dependent. Predator effects were smaller but significant (χ^2^ =5.61, *P*=0.017), leading to an average reduction of around 15–20% in nymph densities.

Alate formation exhibited strong responses to both abiotic and biotic factors, with significant main and interaction effects (*day*: χ^2^ =160.63, P < 0.001; *water × predator*: χ^2^ =65.30, *P*<0.001; *density × predator*: χ^2^ =64.29, *P*<0.001; Fig. 8d). Under well-watered, high-aphid density conditions, predator presence reduced alate numbers by about 55%, whereas under drought, predators increased alate density by over 100%, suggesting an induced dispersal response under combined initial aphid density and drought stress.

## 4. Discussion

Drought is increasingly recognized as a disruptor of trophic interactions, but the mechanisms weakening top-down regulation are not well characterized (Reinecke et al., 2024); here we tested whether drought alters predator-mediated suppression via changes in chemical information that predators use to locate and evaluate high and low prey density patches. Drought reduced plant physiological function and growth, reshaped volatile emission blends, lowered predator visitation, and sharply reduced their reproduction on drought-stressed plants. Under well-watered conditions with high prey density, predators suppressed aphid population growth, suggesting density-mediated top-down effects, and also reshaped aphid demography. Under drought, aphid population growth was constrained by bottom-up limitation, reducing density-mediated predator effects and causing predator influences to be expressed primarily through changes in aphid demographic structure. Thus, drought weakens top-down regulation by disrupting volatile signaling and reducing cues predators use to locate prey.

### Volatile emissions shift with water availability and herbivore density, with consequences for information transfer

Multivariate and univariate compound analyses indicated that volatile composition differed among treatments and that water stress interacted with herbivore density, implying that drought changes both the proportional abundance and functional identity of plant volatile blends (Lin et al., 2022; Rahman et al., 2025). Compound such as methyl salicylate, which is known to increase predator attraction and oviposition (Ayelo, Yusuf, et al., 2021; Russavage et al., 2024; Salamanca et al., 2017; Zhu & Park, 2005), was suppressed under drought, particularly when aphid density was low. For instance, drought suppressed emission of predator-attracting volatiles like methyl salicylate and (E,E)-4,8,12-trimethyltrideca-1,3,7,11-tetraene in tomato, weakening parasitoid attraction and disrupting VOC-mediated indirect defenses (Lin et al., 2022). Similarly, compounds such as (E,E)-α-farnesene and α-terpineol in aphid-infested plants (Truong et al., 2014), and methyl salicylate and (E)-β-ocimene in cassava infested by mites (Pinto-Zevallos et al., 2018), were only significantly emitted at high herbivore densities. This density-dependent induction suggests that at lower infestation levels, VOC blends may be too weak or inconsistent to attract natural enemies. More than just reduced volatile output, shift in blend composition, i.e., altered ratios and compound identities, may render the odor profile less recognizable for natural enemies (Turlings & Erb, 2018), undermining predator attraction and weakening indirect defenses even under high density prey.

### Predators integrate compound-level information, and different compounds may matter for attraction versus commitment

Our findings show that natural enemies do not respond to plant VOCs as a uniform signal but instead exhibit selective, compound-specific responses, consistent with the idea that predators parse volatile blends into functionally distinct cues (Salamanca et al., 2017). Ladybeetles consistently preferred plant VOC profiles associated with well-watered, high-prey density plants, and this preference aligned with elevated levels of methyl salicylate and myrcene. Notably, methyl salicylate was associated with both visitation and oviposition and was reduced under drought, providing a mechanistic link between abiotic stress, altered blend composition, and reduced predator commitment. Importantly, these relationships were identified using VOC profiles measured following sustained herbivory, suggesting that temporally integrated volatile signals are sufficient to capture the biologically relevant odor environment underlying predator decision-making. In contrast, several compounds that were negatively associated with predator responses, including α-pinene, 1-octanol, cryptone, and cymene-7-ol, para, illustrate how deterrent volatiles can shape behavioral outcomes. For instance, Yu et al. (2018) reported that α-pinene reduced prey-searching behavior in insect predators, reinforcing its role as a context-dependent deterrent. These compounds did not necessarily increase in absolute abundance under drought, but shifts in their ratio relative to key candidate attractants may be sufficient to reduce overall blend attractiveness.

This underscores the importance of blend composition and proportionality, as predator responses may depend on a hierarchical sensory evaluation, where certain volatiles function as attractant and others act as deterrents within a blend, making even subtle shifts in their ratios critical to how VOC signals are perceived (Lin et al., 2022; Riddick, 2020; Verheggen et al., 2008). We also observed stage specificity in cue use: for instance, carvacrol was neutral for initial foraging choice but positively predicted oviposition, suggesting that some volatile compound may function as post-settlement cues that reinforce or modulate initial patch assessments (Ayelo, Yusuf, et al., 2021; Verheggen et al., 2008).

Together, these results suggest that predator responses to plant volatiles may be shaped by blend identity and contextual reliability rather than on VOCs presence alone. Such behavior likely reflects evolved decision rules that help predators avoid ovipositing in low-quality or risky patches in heterogeneous environments (Ayelo, Pirk, et al., 2021). Under drought, shift in blend composition, particularly altered ratios and reduced attractant volatiles, may increase signal ambiguity, leading to a higher risks of inflated false negative. As a result, predators may underexploit suitable prey patches when volatile cues no longer exceed the threshold needed to signal patch quality (Tariq et al., 2013).

### Oviposition reveals a stronger threshold response than visitation, and it changes over time

Ladybeetles consistently laid more eggs on well-watered plants with high prey density. In contrast, oviposition on drought-stressed plants, even at high prey density, was rare and only occurred once egg numbers on other treatments were high. This decoupling of prey abundance from predator reproductive investment suggests a behavioral threshold: unless plant VOC blends cross a certain attractant threshold, shaped by their chemical composition and modulated by contextual cues such as prey density and plant water status, predators do not commit to a patch (Verheggen et al., 2008; Xiu et al., 2019). While olfactory cues alone can trigger physiological readiness, such as reduced oosorption and oocyte maturation observed in *Harmonia axyridis* exposed to aphid-induced volatiles even without prey (Rondoni et al., 2017), actual oviposition likely requires the convergence of plant VOCs with other ecological signals that confirm patch reliability.

Importantly, these decisions were temporally structured. Early in the experiment (72-88 hours), oviposition was highly selective and tightly aligned with the most attractive blend. By 110 hours, selectivity weakened, and eggs appeared more often on initially less preferred plants, indicating that patch evaluation is dynamic rather than fixed (Singh et al., 2019). Future studies should extend VOC sampling across the oviposition window to determine whether shifts in blend identity and ratios correlated with this behavioral transition and to identify the volatile cues that maintain commitment over time. In parallel, as egg loads and predator activity accumulated on the most preferred patches, further oviposition there may have increased risks of egg cannibalism or conspecific predation, potentially favoring a broader distribution of eggs across plants to reduce offspring loss (Singh et al., 2019).

Notably, drought-stressed plants often received fewer or no eggs even at high aphid density, suggesting that prey availability cannot fully compensate for lack of reliable chemical signaling. This highlights a critical asymmetry: while herbivore pressure may amplify plant VOC output in some cases, if drought suppresses the specific attractants necessary for decision-making, predators may overlook resource-rich habitats. These results suggest that predator retention and reproductive investment are governed by the convergence of prey abundance, blend identity, and offspring risk, and drought disrupts this convergence by altering VOC blends in ways that suppress predator commitment, even in prey rich patches.

### Bottom-up limitation constrains the expression of top-down control

Drought appears to limit top-down regulation by jointly constraining aphid population growth and weakening attractant cues for predators. Under water stress, aphid abundance was primarily shaped by bottom-up limitation, which reduced the potential for predator to exert additional top-down control, even when predators were present (Kansman et al., 2022). Simultaneously, when water stress altered plant volatile emissions, natural enemies were less likely to invest in foraging and oviposition in prey rich patches (Rahman et al., 2025). In addition to altering prey abundance and signaling, drought may also reduce aphid quality, through changes in size, nutritional content, or chemical composition (Quandahor et al., 2022; Subedi & Kersch-Becker, 2025), which can further discourage predator foraging or reproductive investment. These findings imply that under drought, weak aphid suppression may not be sufficient evidence of weak predator influence because water stress can limit prey population growth and its nutritional quality, and disrupt the chemical cues that structure predator foraging behavior.

While these mechanisms shaped overall aphid suppression, their effects also extended to aphid phenotypic responses, particularly the production of dispersive morphs. However, interpretation of these effects requires caution given the experiment design. Predator presence increased alates on plants under drought but reduced alates under well-watered, high-aphid density conditions, a pattern consistent with predation risk cues shifting aphids toward dispersal phenotypes, including increased production of winged morphs (Hermann et al., 2021). At the same time, because all plant treatments occurred within the same cage and aphids could move among plants, alate counts may also reflect within cage redistribution and aggregation, especially because crowding is itself a potent driver of wing induction (Yuan et al., 2025). As plant VOC blends drew predators to well-watered plants and increased residence time there, predator activity could have displaced aphids toward less visited plants or increased local crowding, either of which can elevate alate production through density-dependent cues (Khallaf et al., 2023).

Future studies that pair repeated VOC sampling with plant specific aphid density and movement tracking will be needed to distinguish developmental induction from redistribution and to test whether odor guided predator attraction and retention drives within cage aggregation patterns.

### Signal contingent trophic regulation under drought and implications

Our results reveal limits to prey-abundance-based models of top-down control by showing that drought undermines trophic cascades not only through bottom-up limitation, but by altering volatile-mediated signaling in ways that reduce predator responsiveness, even when prey are present. We show that predators exhibit compound-specific and behaviorally distinct responses to plant volatiles: methyl salicylate and myrcene increased the likelihood of visitation, while methyl salicylate, n-heptanal, p-cymene, and carvacrol predicted oviposition. Drought suppressed these compounds and shifted blend composition, reducing predator foraging and reproductive investment. As a result, top-down effects shifted from direct consumption to non-consumptive pathways, including changes in aphid alate production and dispersal traits. These findings introduce a signal-contingent framework for understanding trophic regulation: effective control depends not only on prey density, but on the reliability and composition of chemical cues that elicit predator foraging and reproductive responses. When chemical cues are degraded or unreliable, predator populations may decline despite prey availability, weakening trophic cascades and reducing the stability of food webs under environmental stress. Integrating volatile signaling and cue thresholds into multi-species interactions networks will improve predictions of trophic cascades under environmental stress.

## DATA ACCESSIBILITY

Data used in this manuscript are available from the Dryad Digital Repository: http://datadryad.org/share/LINK_NOT_FOR_PUBLICATION/dHwDwkhPMGtfxhuU5vp8DIgT 9Fep1pwDTvesa0LkfgE

## STATEMENT OF AUTHORSHIP

B. Subedi and M.F.K. Becker conceived the study and designed the methodology.

B. Subedi conducted the greenhouse experiments, collected and analyzed the data.

B. Subedi drafted the manuscript. J.G.A and M.F.K. Becker contributed substantially to manuscript writing and interpretation.

All authors critically revised the manuscript and approved the final version for publication.

All authors agree to be accountable for the accuracy and integrity of the work and have met the authorship criteria.

## ACKNOWLEDGEMENTS

We thank Abby Seltzer for her invaluable assistance throughout the course of this study, particularly for her support with data collection, insect rearing, and greenhouse logistics. We also thank Nate McCartney for his help with volatile sample processing and data extractions. This research was supported by the Department of Entomology at the Pennsylvania State University, the National Science Foundation, Division of Environmental Biology award (#2440876), and by the Pennsylvania Department of Agriculture (#C940001869).

