## Supplemental Table 1 for "Drought disrupts volatile-mediated predator foraging and oviposition, weakening trait-mediated top-down control"

**SUPPLEMENTARY INFORMATION**

**Supplementary Table S1** Significant volatile organic compounds (VOCs) correlated with sample separation in NMDS ordination based on Bray-Curtis dissimilarity. Compounds were fitted as vectors onto the NMDS ordination using the *envfit*() function in the vegan R package (999 permutations). The table lists compounds whose correlations with the NMDS ordination were statistically significant (*P*<0.05). The direction (NMDS1 and NMDS2) represents the orientation of each compound vector in ordination space, while R^2^ values indicate the strength of the association between each compound and the overall NMDS structure. Compounds with higher R^2^ values contribute more strongly to sample positioning in NMDS space and are considered important drivers of multivariate chemical variation among treatments

| **Compound** | **CAP1** | **CAP2** | **R2** | **P** | **P-adjusted** |
| --- | --- | --- | --- | --- | --- |
| Hexanal<n-> | -0.931 | 0.365 | 0.364 | 0.001 | 0.0069 |
| Unknown 1 (RI 860.7) | -0.930 | 0.368 | 0.312 | 0.001 | 0.0069 |
| (E)-2-hexenal | -0.570 | -0.821 | 0.312 | 0.001 | 0.0069 |
| Indanol<5-> | -0.598 | 0.802 | 0.288 | 0.001 | 0.0069 |
| Cryptone | -0.823 | -0.568 | 0.277 | 0.001 | 0.0069 |
| heptenal<2E-> | 0.621 | 0.784 | 0.231 | 0.001 | 0.0069 |
| n-Heptanal | -0.992 | 0.129 | 0.203 | 0.001 | 0.0069 |
| Cymen-7-ol<para-> | -0.650 | 0.760 | 0.189 | 0.001 | 0.0069 |
| Decanal<n-> | -0.954 | 0.299 | 0.179 | 0.001 | 0.0069 |
| Furfural | 0.607 | 0.795 | 0.178 | 0.001 | 0.0069 |
| Caryophyllene(E-) | 0.752 | -0.659 | 0.174 | 0.001 | 0.0069 |
| Butyl acetate | 0.891 | -0.454 | 0.172 | 0.001 | 0.0069 |
| Octane, 4-methyl- | -0.853 | 0.521 | 0.171 | 0.001 | 0.0069 |
| 1-Hexanol | -0.982 | -0.188 | 0.153 | 0.001 | 0.0069 |
| hexanal, 2-ethyl- | -0.990 | -0.141 | 0.143 | 0.001 | 0.0069 |
| Pentanol<3-methyl-> | -0.782 | 0.624 | 0.141 | 0.001 | 0.0069 |
| Undecenol<2E-> | -0.997 | 0.082 | 0.109 | 0.001 | 0.0069 |
| Lavandulyl<tetrahydro-> | -0.551 | 0.835 | 0.138 | 0.002 | 0.0107 |
| Octanol<n-> | -0.974 | -0.226 | 0.129 | 0.002 | 0.0107 |
| Nonanal<n-> | -0.955 | 0.296 | 0.123 | 0.002 | 0.0107 |
| Tridecenol<2E-> | -0.930 | 0.366 | 0.102 | 0.002 | 0.0107 |
| Dihydro citronellol acetate | 0.812 | -0.584 | 0.086 | 0.002 | 0.0107 |
| Sabinene | -0.872 | 0.490 | 0.131 | 0.003 | 0.0141 |
| Carvacrol | 0.828 | -0.561 | 0.108 | 0.003 | 0.0141 |
| Hexenyl angelate<3Z-> | -0.825 | -0.565 | 0.087 | 0.003 | 0.0141 |
| Hexenyl acetate<3E-> | -0.563 | 0.826 | 0.122 | 0.004 | 0.017 |
| Acetylacetophenone<para- | -0.964 | 0.264 | 0.103 | 0.004 | 0.017 |
| alpha-phellandrene | -0.124 | -0.992 | 0.090 | 0.006 | 0.0236 |
| Cresol acetate<para- | -0.360 | -0.933 | 0.084 | 0.006 | 0.0236 |
| Hexenyl butanoate<3Z-> | -0.810 | 0.586 | 0.083 | 0.006 | 0.0236 |
| Isoborneol<8-isobutyryloxy-> | -0.966 | -0.257 | 0.082 | 0.007 | 0.025 |
| 3-methyl-1-butanol | 0.959 | -0.282 | 0.081 | 0.007 | 0.025 |
| Thujene<alpha-> | 0.998 | 0.060 | 0.093 | 0.008 | 0.027 |
| beta-phellandrene | 0.608 | -0.794 | 0.088 | 0.008 | 0.027 |
| (Z)-3-hexenol | -0.634 | 0.774 | 0.090 | 0.01 | 0.031 |
| 5-Hepten-2-one, 6-methyl- | -0.991 | 0.133 | 0.087 | 0.01 | 0.031 |
| 2-methyl-1-butanol | -0.819 | -0.574 | 0.077 | 0.01 | 0.031 |

**Supplementary Table S2** Estimated marginal means (± SE) for volatile organic compound (VOC) emissions under water treatment, aphid density, and Water × Density interaction models. The table combines outputs from the water-only, density-only, and interaction analyses for all compounds. Different letters indicate significant differences among treatment levels based on post hoc comparisons.

| **Compound** | **Water** | **Density** | **Mean** | **SE** | **Significnace Letter** |
| --- | --- | --- | --- | --- | --- |
| Santalol acetate<(Z)-epi-beta-> | Well-watered | Low | 0.122 | 0.015 | a |
|  | Well-watered | High | 0.110 | 0.014 | a |
|  | Drought | Low | 0.162 | 0.020 | a |
|  | Drought | High | 0.130 | 0.016 | a |
| Sabinene | Well-watered | Low | 0.623 | 0.111 | ab |
|  | Well-watered | High | 0.802 | 0.140 | b |
|  | Drought | Low | 0.590 | 0.105 | ab |
|  | Drought | High | 0.389 | 0.075 | a |
| Pinocampheol | Well-watered | Low | 0.021 | 0.004 | b |
|  | Well-watered | High | 0.026 | 0.007 | b |
|  | Drought | Low | 0.010 | 0.002 | a |
|  | Drought | High | 0.014 | 0.002 | ab |
| Penten-1-al<2E-> | Well-watered | Low | 0.006 | 0.002 | ab |
|  | Well-watered | High | 0.005 | 0.001 | ab |
|  | Drought | Low | 0.003 | 0.001 | a |
|  | Drought | High | 0.012 | 0.005 | b |
| Pentanol<3-methyl-> | Well-watered | Low | 0.013 | 0.003 | b |
|  | Well-watered | High | 0.004 | 0.001 | a |
|  | Drought | Low | 0.004 | 0.001 | a |
|  | Drought | High | 0.007 | 0.002 | ab |
| p-cymene | Well-watered | Low | 0.042 | 0.006 | a |
|  | Well-watered | High | 0.060 | 0.008 | a |
|  | Drought | Low | 0.054 | 0.008 | a |
|  | Drought | High | 0.044 | 0.006 | a |
| Octen-3-one<1-> | Well-watered | Low | 0.016 | 0.006 | c |
|  | Well-watered | High | 0.014 | 0.005 | b |
|  | Drought | Low | 0.012 | 0.004 | a |
|  | Drought | High | 0.039 |  |  |
| Octane, 4-methyl- | Well-watered | Low | 0.047 | 0.005 | a |
|  | Well-watered | High | 0.046 | 0.005 | a |
|  | Drought | Low | 0.052 | 0.005 | a |
|  | Drought | High | 0.053 | 0.005 | a |
| Ocimene<(Z)-beta-> | Well-watered | Low | 0.091 | 0.018 | a |
|  | Well-watered | High | 0.085 | 0.013 | a |
|  | Drought | Low | 0.055 | 0.011 | a |
|  | Drought | High | 0.054 | 0.010 | a |
| Nonanal<n-> | Well-watered | Low | 0.112 | 0.011 | a |
|  | Well-watered | High | 0.103 | 0.010 | a |
|  | Drought | Low | 0.139 | 0.013 | a |
|  | Drought | High | 0.124 | 0.012 | a |
| Methyl salicylate | Well-watered | Low | 0.020 | 0.004 | a |
|  | Well-watered | High | 0.039 | 0.007 | b |
|  | Drought | Low | 0.018 | 0.004 | a |
|  | Drought | High | 0.031 | 0.005 | ab |
| Longicyclene | Well-watered | Low | 0.046 | 0.003 | a |
|  | Well-watered | High | 0.074 | 0.004 | b |
|  | Drought | Low | 0.065 | 0.005 | b |
|  | Drought | High | 0.014 |  |  |
| Lavandulyl<tetrahydro-> | Well-watered | Low | 0.018 | 0.006 | b |
|  | Well-watered | High | 0.005 | 0.002 | a |
|  | Drought | Low | 0.015 | 0.005 | b |
|  | Drought | High | 0.009 | 0.003 | ab |
| Indanol<5-> | Well-watered | Low | 0.040 | 0.005 | a |
|  | Well-watered | High | 0.063 | 0.016 | a |
|  | Drought | Low | 0.032 | 0.003 | a |
|  | Drought | High | 0.036 | 0.005 | a |
| heptane, 4-methyl- | Well-watered | Low | 0.007 | 0.001 | ab |
|  | Well-watered | High | 0.006 | 0.001 | a |
|  | Drought | Low | 0.008 | 0.001 | b |
|  | Drought | High | 0.007 | 0.001 | ab |
| Heptanal | Well-watered | Low | 0.007 | 0.001 | ab |
|  | Well-watered | High | 0.008 | 0.001 | ab |
|  | Drought | Low | 0.012 | 0.002 | b |
|  | Drought | High | 0.006 | 0.001 | a |
| Dodecene<1-> | Well-watered | Low | 0.085 | 0.021 | b |
|  | Well-watered | High | 0.073 | 0.020 | b |
|  | Drought | Low | 0.067 | 0.014 | b |
|  | Drought | High | 0.025 | 0.006 | a |
| Dihydroisojasmone | Well-watered | Low | 0.046 | 0.007 | a |
|  | Well-watered | High | 0.046 | 0.008 | a |
|  | Drought | Low | 0.031 | 0.005 | a |
|  | Drought | High | 0.037 | 0.006 | a |
| Decane<n-> | Well-watered | Low | 0.005 | 0.001 | a |
|  | Well-watered | High | 0.005 | 0.001 | a |
|  | Drought | Low | 0.006 | 0.001 | a |
|  | Drought | High | 0.008 | 0.001 | a |
| Cumene | Well-watered | Low | 0.021 | 0.002 | ab |
|  | Well-watered | High | 0.020 | 0.002 | a |
|  | Drought | Low | 0.025 | 0.002 | bc |
|  | Drought | High | 0.027 | 0.002 | c |
| Butyl acetate | Well-watered | Low | 0.002 | 0.000 | ab |
|  | Well-watered | High | 0.002 | 0.000 | a |
|  | Drought | Low | 0.002 | 0.000 | ab |
|  | Drought | High | 0.002 | 0.000 | b |
| Benzeneacetaldehyde | Well-watered | Low | 0.011 | 0.003 | b |
|  | Well-watered | High | 0.006 | 0.002 | a |
|  | Drought | Low | 0.019 | 0.005 | c |
|  | Drought | High | 0.009 | 0.002 | b |
| 2(5H)-Furanone | Well-watered | Low | 0.002 | 0.002 | ab |
|  | Well-watered | High | 0.005 | 0.003 | bd |
|  | Drought | Low | 0.003 | 0.002 | cd |
|  | Drought | High | 0.004 | 0.002 | ac |
| Unknown 1 (RI 860.7) | Well-watered | Low | 0.003 | 0.001 | a |
|  | Well-watered | High | 0.001 | 0.001 | a |
|  | Drought | Low | 0.005 | 0.001 | a |
|  | Drought | High | 0.005 | 0.001 | a |
